# CD47 suppresses phagocytosis by repositioning SIRPA and preventing integrin activation

**DOI:** 10.1101/752311

**Authors:** Meghan A. Morrissey, Ronald D. Vale

## Abstract

Macrophages must engulf dead cells, debris, and pathogens, while selecting against healthy cells to prevent autoimmunity. Healthy cells express CD47 on their surface, which activates the SIRPA receptor on macrophages to suppress engulfment. Cancer cells overexpress CD47 to evade clearance by the innate immune system, making the CD47-SIRPA signaling axis an appealing therapeutic target. However, the mechanism by which CD47-SIRPA inhibits engulfment remains poorly understood. Here, we dissect SIRPA signaling using a reconstituted target with varying concentrations of activating and inhibitor ligands. We find that SIRPA is excluded from the phagocytic synapse between the macrophage and its target unless CD47 is present. Artificially directing SIRPA to the kinase-rich synapse in the absence of CD47 activates SIRPA and suppresses engulfment, indicating that the localization of the receptor is critical for inhibitory signaling. CD47-SIRPA inhibits integrin activation in the macrophage, reducing macrophage-target contact and suppressing phagocytosis. Chemical activation of integrins can override this effect and drive engulfment of CD47-positive targets, including cancer cells. These results suggest new strategies for overcoming CD47-SIRPA inhibition of phagocytosis with potential applications in cancer immunotherapy.

## Introduction

The innate immune system is finely balanced to rapidly activate in response to pathogenic stimuli, but remain quiescent in healthy tissue. Macrophages, key effectors of the innate immune system, measure activating and inhibitory signals to set a threshold for engulfment and cytokine secretion. The cell surface protein CD47 is a “Don’t Eat Me” signal that protects healthy cells from macrophage engulfment and is often upregulated by cancer cells to evade innate immune detection (Chao et al., 2012; Jaiswal et al., 2009; Majeti et al., 2009; Oldenborg et al., 2001, 2000). CD47 function-blocking antibodies result in decreased cancer growth or tumor elimination (Advani et al., 2018; Chao et al., 2010a; Gholamin et al., 2017; Jaiswal et al., 2009; Willingham et al., 2012). Despite the therapeutic promise of manipulating CD47 signaling, the mechanism by which CD47 suppresses macrophage signaling is unclear.

CD47 on the surface of cancer cells binds to SIRPA in macrophages or dendritic cells to prevent activation (Jiang et al., 1999; Liu et al., 2015; Okazawa et al., 2005; Seiffert et al., 1999; Tseng et al., 2013; Yi et al., 2015). Activation of the inhibitory receptor SIRPA must be controlled with high fidelity to suppress engulfment of viable cells when CD47 is present while allowing for robust engulfment of targets lacking CD47. CD47 binding triggers SIRPA phosphorylation by Src family kinases (Barclay and Brown, 2006), but how CD47 binding is translated across the cell membrane to drive SIRPA phosphorylation is not known. Activated SIRPA recruits the phosphatases SHP-1 and SHP-2 (Fujioka et al., 1996; Noguchi et al., 1996; Okazawa et al., 2005; Oldenborg et al., 2001; Veillette et al., 1998). The downstream events that shut off the engulfment program are not clear.

In vivo, CD47 has been reported to suppress multiple different pro-engulfment “Eat Me” signals, including IgG, complement and calreticulin (Chen et al., 2017; Gardai et al., 2005; Oldenborg et al., 2001). This complexity, in addition to substantial variation in target size, shape and concentration of “Eat Me” signals, can make a quantitative, biochemical understanding of receptor activation difficult. To overcome this problem, we utilize a synthetic target cell-mimic with a defined complement of signals to interrogate the mechanism of SIRPA activation and its downstream targets. We find that CD47 ligation alters SIRPA localization, positioning SIRPA for activation at the phagocytic synapse. At the phagocytic synapse, SIRPA inhibits integrin activation to limit macrophage spreading across the surface of the engulfment target. Directly activating integrin eliminated the effect of CD47 and rescued engulfment. Activation of integrin also allowed macrophages to engulf cancer cells, similar to the effect observed with a CD47 function-blocking antibody.

## Results

### CD47-SIRPA signaling suppresses IgG and phosphatidylserine “Eat Me” signals

To study the mechanism of “Eat Me” and “Don’t Eat Me” signal integration during engulfment, we used a reconstituted engulfment target (Figure 1A). Silica beads were coated in a supported lipid bilayer to mimic the surface of a cancer cell. To activate engulfment, we introduced IgG, a well-defined “Eat Me” signal that synergizes with CD47 blockade to promote cancer cell clearance (Chao et al., 2010a; Freeman and Grinstein, 2014). IgG is recognized by the Fc γ Receptor family (FcR), which activates downstream signaling and engulfment (Freeman and Grinstein, 2014). To activate SIRPA, we incorporated the CD47 extracellular domain at a surface density selected to mimic the CD47 density on cancer cells (∼600 molecules/μm^2^, Figure S1).

**Figure 1:**
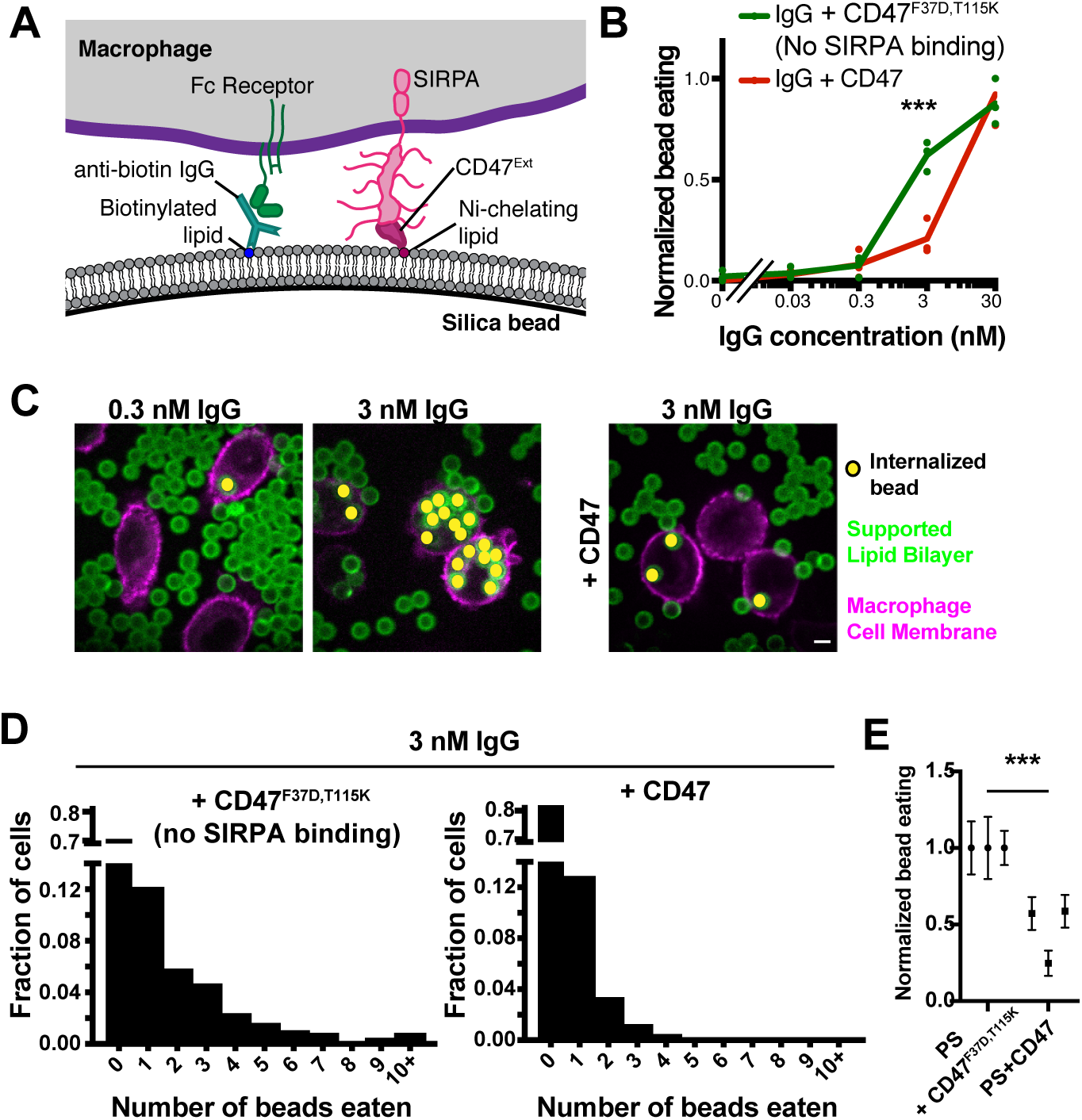
CD47-SIRPA suppresses IgG and PS dependent engulfment. (A) Schematic shows the supported lipid bilayer system used in this study. Anti-biotin IgG is bound to biotinylated lipids. IgG is recognized by Fc Receptor in the macrophage. The extracellular domain of CD47-His_10_ is bound to Ni-NTA-conjugated lipids and recognized by SIRPA in the macrophage. (B) Silica beads are coated in a supported lipid bilayer and incubated with the indicated concentration of IgG and either CD47 (red) or an inactive mutant CD47 (F37D, T115K; green). The functionalized beads were added to RAW264.7 macrophages and fixed after 30 min. The average number of beads per macrophage was assessed by confocal microscopy and normalized to the maximum bead eating observed in that replicate. Each dot represents an independent replicate (n≥100 cells analyzed per experiment), and the trendline connects the average of three replicates. (C) Still images depict the assay described in (B). The supported lipid bilayers contain the fluorescently-labeled lipid atto390-DOPE (green) and the macrophages membranes are labeled with CellMask (magenta). Internalized beads are indicated with a yellow dot. (D) Histograms depict the fraction of cells engulfing the indicated number of beads (pooled data from the three independent replicates shown in (B)). Macrophages encountering CD47-conjugated beads (right) were less likely to engulf, and those that did engulfed fewer beads. CD47^F37D,T115K^, a mutant that cannot bind SIRPA, was used as a control. (E) Macrophages were incubated with beads coated in a supported lipid bilayer containing 10% phosphatidylserine and either CD47 or the inactive CD47^F37D,T115K^. Data was normalized to the maximum bead eating observed in that replicate. The complete, pooled data is shown in Supplementary Figure 1E. Dots and error bars denote the mean and standard error of independent replicates. * * * indicates p<0.0005 by a Kruskal-Wallis test on the pooled data (B and E). Scale bar denotes 5 µm in this and all subsequent figures.

Using this system, we tested the effect of CD47 on engulfment across a titration of IgG densities (Figure S1). We mixed beads with the macrophage-like cell line RAW264.7 and measured the number of internalized beads by confocal microscopy. We found that CD47-SIRPA signaling suppressed engulfment at intermediate IgG densities, but did not appreciably affect engulfment of targets with high densities of bound IgG (Figure 1B-D). This suppression was dependent on CD47 binding as a mutated CD47 extracellular domain (F37D, T115K) that is unable to bind to SIRPA (Hatherley et al., 2008) was also unable to suppress engulfment.

We further examined whether CD47-SIRPA signaling could suppress engulfment of targets mimicking apoptotic corpses. A critical “Eat Me” signal from apoptotic corpses is phosphatidylserine, which becomes exposed on the outer leaflet of the plasma membrane during cell stress, apoptosis (Fadok et al., 1992; Poon et al., 2014), and on some cancer cells (Birge et al., 2016; Utsugi et al., 1991). We found that engulfment of beads containing 10% phosphatidylserine in the supported lipid bilayer was inhibited by the inclusion of CD47 on the bilayer (Figure 1E, Figure S1). Together, these data show that CD47-SIRPA signaling can block engulfment driven by IgG and phosphatidylserine. Thus, bilayer-coated beads provide a well-defined and tunable platform for studying the integration of “Eat Me” and “Don’t Eat Me” signals during engulfment.

### CD47 ligation relocalizes SIRPA to the phagocytic synapse

We next sought to determine the mechanism by which CD47 ligation regulates SIRPA activity. We first examined SIRPA localization during phagocytosis of IgG-coated beads. In the absence of CD47, SIRPA was segregated away from the phagocytic cup that enveloped IgG-coated beads (Figure 2A). Similarly, SIRPA was depleted at the center of the immunological synapse between a macrophage and a supported lipid bilayer containing phosphatidylserine (Figure S2). In contrast, in the presence of CD47, SIRPA remained at the phagocytic cup (Figure 2A). These data demonstrate that CD47 recruits SIRPA to the phagocytic synapse.

**Figure 2:**
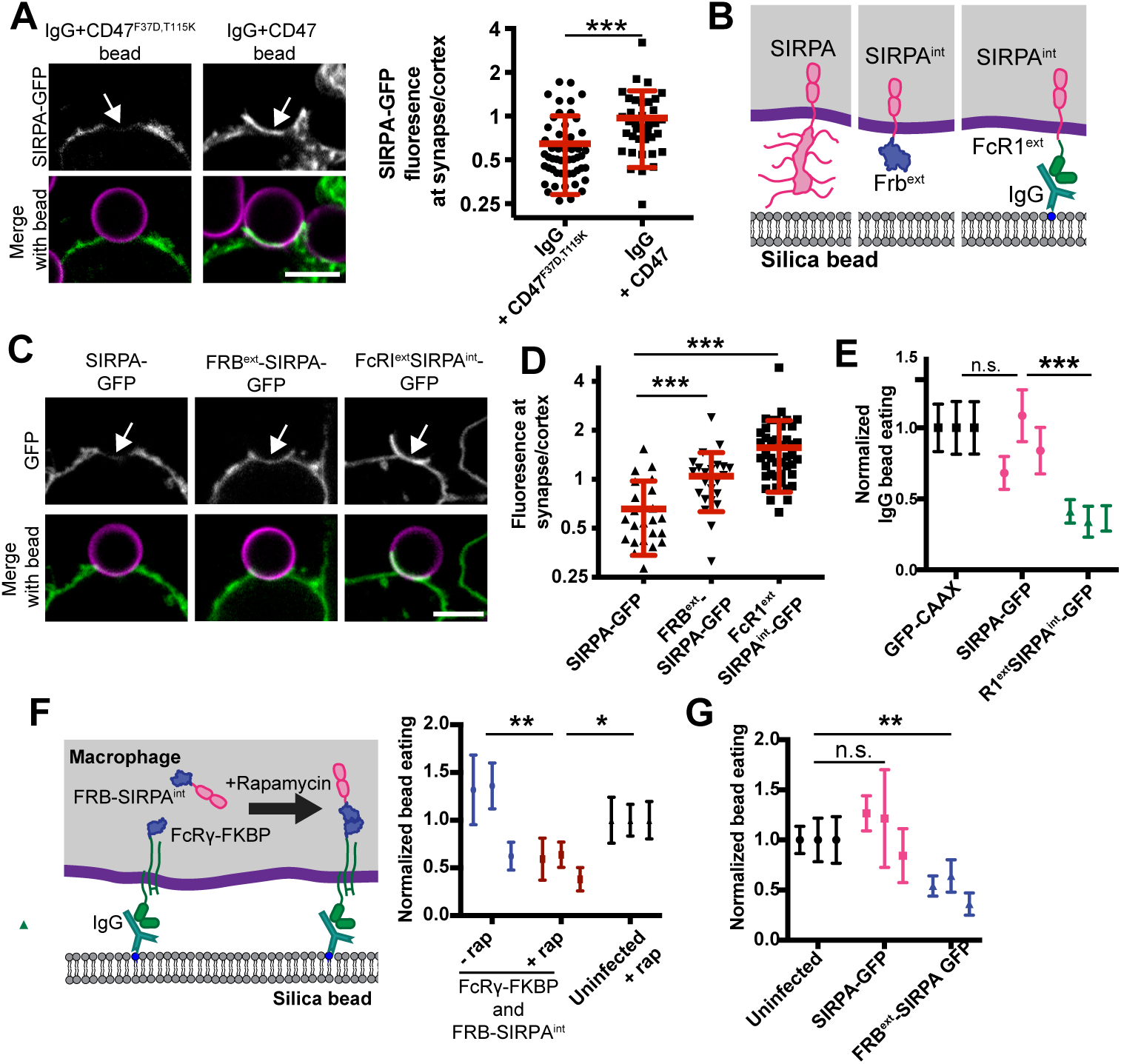
Forcing SIRPA into the macrophage-target synapse suppresses engulfment. (A) SIRPA-GFP (top; green in merge) is depleted from the base of the phagocytic cup (arrow) when a macrophage engulfs a bead functionalized with IgG and CD47^F37D,^ T115K, which cannot bind SIRPA (left; supported lipid bilayer, magenta). SIRPA is not depleted when CD47 is present (IgG+CD47, right). Graph depicts the ratio of SIRPA-GFP at the phagocytic cup/cell cortex for individual phagocytic cups. (B) A schematic shows the chimeric SIRPA constructs in this figure. Full length SIRPA is on the right, FRB^ext^-SIRPA is in the center and FcRI^ext^-SIRPA^int^-GFP is on the left. (C) SIRPA-GFP, FRB^ext^-SIRPA-GFP and FcRI^ext^-SIRPA^int^-GFP fluorescence is shown at cell-bead contacts (arrow). (D) Graph depicts the ratio of GFP fluorescence at the synapse (arrow in C) compared to the cortex for the indicated SIRPA chimeras. (E) A graph depicts the average number of internalized IgG beads per macrophage expressing the chimeric SIRPA constructs schematized in (B), normalized to macrophages expressing only a membrane-tethered GFP (GFP-CAAX). (F) Schematic (left) shows a system for inducible recruitment of the SIRPA intracellular domain to the phagocytic cup. Recruiting SIRPA to the phagocytic cup suppresses engulfment compared to soluble SIRPA or compared to wild-type macrophages treated with rapamycin (normalized to uninfected macrophages). (G) The graph shows the number of beads engulfed by uninfected, SIRPA-GFP or FRB^ext^-SIRPA expressing macrophages normalized to uninfected cells. In A and D, dots represent individual cups, red lines show mean ± SD and data is pooled from three independent experiments. In E, F and G, dots show the average from an independent replicate with the error bars denoting SEM for that replicate. The complete pooled data showing the number of beads eaten per macrophage is shown in Figure S2. * * * denotes p<0.0005, * * denotes p<0.005 and * denotes p<0.05 as determined by a Student’s T test (A, D) or a Kruskal-Wallis test on the pooled data from all three replicates (E, F, G).

We next sought to address the mechanism of SIRPA segregation away from the phagocytic cup in the presence of IgG and absence of CD47. We hypothesized that exclusion of unligated SIRPA from the synapse could be driven by its heavily glycosylated extracellular domain, either by interactions with the surrounding glycocalyx or steric exclusion from the spatially restricted phagocytic synapse. We therefore created a SIRPA chimeric receptor where the extracellular domain was replaced with a small, inert protein domain (FRB^ext^-SIRPA; Figure 2B). Unlike full length SIRPA, FRB^ext^-SIRPA was not segregated away from the cell-target synapse (Figure 2C, D). This result demonstrates that the extracellular domain of SIRPA is required for SIRPA exclusion from the phagocytic cup.

### Targeting SIRPA to the phagocytic synapse suppresses engulfment

Receptor activation by Src family kinases at the phagocytic cup is favored due to exclusion of bulky phosphatases like CD45 (Freeman et al., 2016; Goodridge et al., 2011). We therefore hypothesized that positioning SIRPA at the phagocytic cup may drive receptor activation. To distinguish between the effects of CD47 binding and synapse localization, we developed a chimeric SIRPA receptor that localized to the phagocytic synapse in the absence of CD47. We replaced the SIRPA extracellular domain with the IgG-binding extracellular domain of the FcγRI α chain (Figure 2B; termed FcR1^ext^-SIRPA^int^). This receptor is driven into the synapse by IgG binding instead of CD47 (Figure 2C, D). Expression of this synapse-localized chimera suppressed engulfment of IgG-coated beads in the absence of CD47 (Figure 2E, Figure S2). Thus, targeting SIRPA to the phagocytic cup is sufficient to inhibit engulfment, even in the absence of its natural ligand CD47.

As an alternative strategy to control the localization of SIRPA activity, we used a chemically inducible dimerization system (Spencer et al., 1993). We fused one half of the chemically inducible dimer to FcR (FcR γ chain-FKBP) and the second to a soluble SIRPA intracellular domain (FRB-SIRPA^int^, Figure 2F). In the presence of the small molecule rapamycin, FKBP and FRB form a high-affinity dimer (Spencer et al., 1993), thereby recruiting the SIRPA intracellular domain to FcR. In the absence of rapamycin, cells efficiently engulfed IgG-coated beads (Figure 2F, Figure S2). In contrast, rapamycin-induced recruitment of the SIRPA intracellular domain to the FcR γ chain significantly suppressed engulfment (Figure 2F).

We next examined the extracellular domain truncation of SIRPA (FRB^ext^-SIRPA) that was not excluded from the phagocytic synapse (Figure 2B). FRB^ext^-SIRPA constitutively suppressed engulfment (Figure 2G), demonstrating that exclusion of SIRPA is essential for efficient engulfment. Taken together, these experiments show that CD47 activates SIRPA by recruiting it to the phagocytic synapse.

### FcR phosphorylation is not a major target of CD47-SIRPA signaling

We next sought to determine how activated SIRPA inhibits engulfment. Phosphorylated SIRPA recruits the phosphatases SHP-1 and SHP-2 via their phosphobinding SH2 domains but the downstream targets of SHP-1 and SHP-2 are not known (Fujioka et al., 1996; Noguchi et al., 1996; Okazawa et al., 2005; Oldenborg et al., 2001; Veillette et al., 1998). One potential target of SIRPA-bound SHP phosphatases is FcR itself. When encountering an IgG-bound bilayer, macrophages clustered IgG into mobile microclusters (Figure 3, Movie S1) that recruited Syk (Figures S3 (Lin et al., 2016). When CD47 was present, these microclusters still formed and recruited Syk, suggesting that FcR is still phosphorylated (Figures 3A and S3, Movie S2). Further, when we looked at SIRPA localization at the cell-target interface at high resolution, we found that, even in the presence of CD47, SIRPA did not co-localize with FcR clusters, suggesting that SIRPA is not positioned to dampen receptor activation (Figure S3). Overall, this suggests that changes to FcR activation and Syk recruitment are unlikely to account for the effect of SIRPA, consistent with previous biochemical observations (Okazawa et al., 2005; Tsai and Discher, 2008).

**Figure 3:**
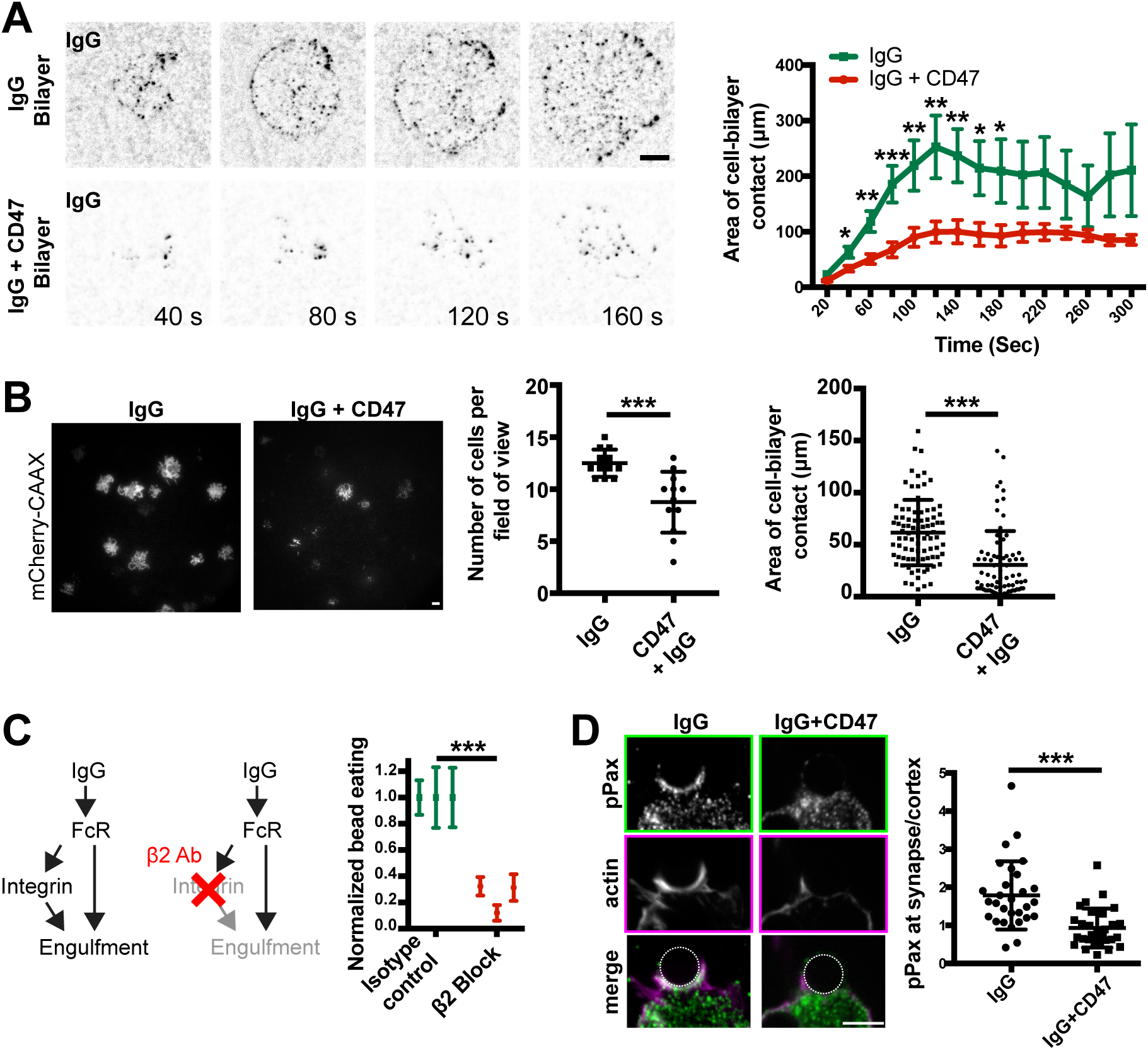
CD47 prevents integrin activation. (A) Still images from a TIRF microscopy timelapse show that macrophages form IgG (black) microclusters as they spread across an IgG bilayer (top). Adding CD47 to the bilayer inhibits cell spreading (bottom; graphed on right, average area of contact from n≥11 cells ± SEM, pooled from three separate experiments). (B) TIRF images show the cell membrane (mCherry-CAAX; white) of macrophages engaging with an IgG (left) or IgG and CD47 (right) bilayer. Graphs depict the average number of cells seen contacting the bilayer after 10 min (center) and the average area of cell contact (right). Each dot represents an individual field of view (center) or cell (right) pooled from three independent experiments. (C) Diagram shows that IgG binding activates Fc Receptor, which triggers downstream signaling events including inside-out activation of integrins. Blocking integrin activation using a function-blocking antibody (2E6) targeting the β2 integrin subunit decreased the efficiency of engulfment (graphed in center panel, normalized to the isotype control, with error bars denoting SEM of each replicate). (D) Immunofluorescence images show phosphopaxillin (top; green in merge) and F-actin (center; magenta in merge; visualized with phalloidin) at the phagocytic cup of an IgG coated bead (left) or an IgG- and CD47-coated bead (right). Graphs show the ratio of phosphopaxilin intensity at the phagocytic cup/cell cortex. Each dot represents an individual phagocytic cup; lines denote the mean ± SD. The non-activating CD47^F37D, T115K^ was used as a control on bilayers lacking CD47. * * * denotes p<0.0005, * * denotes p<0.005, and * denotes p<0.05 as determined by Student’s T test (A, B and D) or a Kruskal-Wallis test on the pooled data (C).

### CD47 prevents integrin activation

During our TIRF experiments, we observed a difference in the spreading of cells on bilayers containing IgG alone versus IgG plus CD47. On IgG-coated bilayers, cells rapidly spread across the bilayer surface (Figure 3A, Movie S1). In contrast, macrophages encountering an IgG and CD47-containing bilayer exhibited reduced cell spreading (Figure 3A and 3B, Movie S2). These data shows that CD47 inhibits cell spreading across a target substrate.

Cell spreading is thought to involve activation of integrins and the actin cytoskeleton (Springer and Dustin, 2011). Inactive integrins exist in a low affinity, bent confirmation (Springer and Dustin, 2011). Upon activation, the extracellular domain extends into an open conformation that can bind many ligands with high affinity (Freeman and Grinstein, 2014; Springer and Dustin, 2011). FcR activation stimulates inside-out activation of integrins (Dupuy and Caron, 2008; Jones et al., 1998). Activated integrins can then promote engulfment, either by increasing adhesion to the target particle or by reorganizing the actin cytoskeleton (Dupuy and Caron, 2008; Wong et al., 2016). We found that inhibiting integrin with a β2 integrin function-blocking antibody (2E6) or Fab dramatically decreased engulfment efficiency (Figure 3C and S3), demonstrating that inactivating integrins is sufficient to suppress engulfment.

Because integrin is required for cell spreading and engulfment (Springer and Dustin, 2011), we hypothesized that CD47-SIRPA signaling may inhibit engulfment by preventing inside-out activation of integrin. Supporting this hypothesis, a previous study identified phosphopaxillin, which is specifically recruited to sites of integrin activation, as one of a number of phosphoproteins affected by CD47 (Geiger et al., 2009; Tsai and Discher, 2008). We found that the enrichment of phospho-paxillin at the interface of the macrophage with an IgG-coated bead was substantially diminished by the simultaneous presence of CD47 on the bead (Figure 3D). Together, these data indicate that CD47-SIRPA signaling prevents integrin activation.

### Activating integrin bypasses CD47-SIRPA inhibitory signaling

CD47-SIRPA has previously been reported to affect paxillin and myosin phosphorylation, as well as F-actin recruitment (Tsai and Discher, 2008). It is not clear which of these is a target of CD47 signaling and which is a secondary effect of altered upstream signaling (Tsai and Discher, 2008). We hypothesized that if SIRPA signaling suppresses engulfment primarily by inhibiting integrin inside-out activation, then directly activating integrin might bypass SIRPA-mediated inhibition and permit bead engulfment (Figure 4A). Alternatively, if the target of CD47-SIRPA signaling is in a parallel pathway or downstream of integrin activation, then activating integrin should not rescue engulfment following SIRPA activation. To activate integrin, we treated macrophages with manganese, which locks integrin into a high-affinity open conformation (Dransfield et al., 1992). We found that macrophages treated with 1 mM manganese engulfed beads with a similar efficiency whether or not CD47 was conjugated to the supported lipid bilayer (Figure 4B). Importantly, manganese did not trigger bead engulfment on its own or dramatically enhance engulfment of IgG-coated beads in the absence of CD47 (Figure 4B,C), establishing that increasing integrin activation is not sufficient to trigger engulfment. Thus, a manganese-induced increase in engulfment was specific to beads coated with CD47 and IgG.

**Figure 4:**
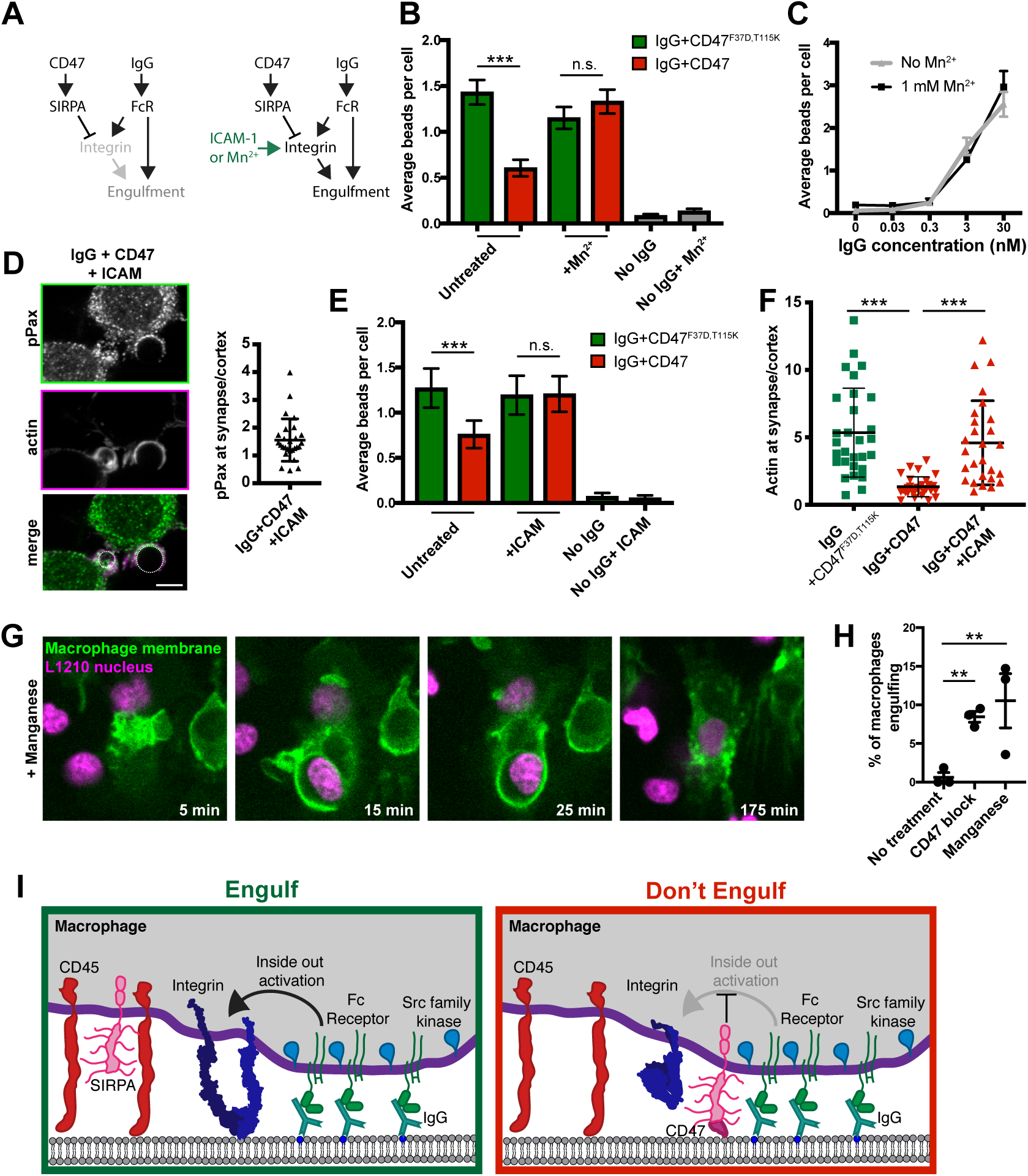
Bypassing inside out activation of integrin eliminates the effect of CD47. (A) The schematic shows a simplified signaling diagram. If CD47 and SIRPA act upstream of integrin, then providing an alternate means of integrin activation (Mn^2+^ or ICAM) should eliminate the effect of CD47. (B) Macrophages were treated with 1 mM Mn^2+^ and fed beads with IgG and either CD47 (red) or the non-signaling CD47^F37D,^ T115K (green). Bars denote the average number of beads eaten from the pooled data of three independent replicates ± SEM. (C) Beads were incubated with the indicated concentration of IgG and added to macrophages. Treatment with Mn^2+^ did not dramatically enhance engulfment (black, compared to grey). Dots represent the average number of beads eaten ± SEM in one data set representative of three experiments. (D) Immunofluorescence shows that adding ICAM (10 nM coupling concentration) to IgG + CD47 beads rescues phosphopaxillin (top; green in merge, bottom) at the phagocytic cup. Compare to data displayed in Figure 3D (p<0.0005 for phosphopaxilin with ICAM and CD47 compared to CD47 alone). (E) Beads were functionalized with IgG and either CD47 (red) or the non-signaling CD47^F37D, T115K^ (green). Adding ICAM to the beads abrogated the effect of CD47 (center) but did not stimulate engulfment without IgG (right). (F) ICAM also rescued actin accumulation at the phagocytic cup as measured by the ratio of phalloidin fluorescence at the cup to the cell cortex. (G) Bone marrow-derived macrophages expressing a membrane tethered GFP (GFP-CAAX) were incubated with L1210 murine leukemia cells expressing H2B-mCherry. Treating with 100 μM manganese allowed for engulfment of whole cancer cells. These images correspond to frames from Movie S3. (H) The percent of macrophages engulfing a cancer cell during an 8 hr timelapse is graphed. Each dot represents an independent replicate, with lines denoting mean ± SEM. (I) Model figure shows that in the absence of CD47 (left), SIRPA is segregated away from the phagocytic synapse and Fc Receptor binding triggers inside out activation of integrin. When CD47 is present (right), SIRPA localizes to the synapse and inhibits integrin activation. * * * denotes p<0.0005, * * denotes p<0.005 and n.s. denotes p>0.05 as determined by a Kruskal-Wallis test (B, E), Ordinary One-way ANOVA (D, F) or Fisher Exact (H) on the pooled data from all three replicates.

As an alternative strategy to activate integrins, we incubated macrophages with beads containing a surplus of high affinity integrin ligand, ICAM-1 (Springer and Dustin, 2011). ICAM-1 was sufficient to activate integrin and recruit phophopaxillin even in the presence of CD47 (Figure 4D). Inclusion of high concentrations of ICAM-1 abrogated the inhibitory effect of CD47 on phagocytosis, but did not dramatically alter the engulfment efficiency of IgG coated beads in the absence of CD47 (Figure 4E).

CD47 has previously been reported to inhibit downstream steps in the phagocytic signaling pathway, including actin accumulation at the phagocytic cup (Tsai and Discher, 2008). Despite the presence of CD47, ICAM-1-bound beads had similar levels of actin accumulation as beads lacking CD47 (Figure 4F). This demonstrates that activating integrins reactivates downstream signaling in the presence of CD47. Together, these data suggest that inside-out activation of integrin is the primary target of CD47-SIRPA signaling.

### Integrin activation drives cancer cell engulfment

Many cancer cells overexpress CD47 to evade the innate immune system despite increased expression of “Eat Me” signals such as calreticulin or phosphatidylserine (Birge et al., 2016; Chao et al., 2010b; Gardai et al., 2005; Utsugi et al., 1991). Blocking CD47 with a therapeutic antibody allows “Eat Me” signals to dominate, resulting in engulfment of whole cancer cells (Jaiswal et al., 2009; Majeti et al., 2009). We hypothesized that exogenous activation of integrin would bypass the CD47 signal on the surface of cancer cells, allowing for engulfment. To test this, we incubated bone marrow derived mouse macrophages with a CD47-positive murine leukemia line, L1210 (Chen et al., 2017). We found that activating integrins with 100 µM manganese increased the ability of macrophages to engulf cancer cells, reaching a similar efficiency as treatment with a CD47 function-blocking antibody (Figure 4G,H; Movie S3). Manganese did not directly affect cancer cell viability over the time course of this experiment (Figure S4). This data shows that activating integrins bypasses the suppressive CD47 signal on the surface of cancer cells.

## Discussion

CD47-SIRPA signaling suppresses engulfment, protecting viable cells and allowing cancer cells to evade the innate immune system (Jaiswal et al., 2009; Majeti et al., 2009; Oldenborg et al., 2000). Although CD47 blockade is a promising new target for cancer therapies (Advani et al., 2018; Gholamin et al., 2017; Willingham et al., 2012), the mechanism of CD47-SIRPA signaling has not been clarified. We demonstrate that localizing SIRPA to the phagocytic synapse is sufficient to activate this inhibitory receptor. Once active, SIRPA suppresses engulfment by preventing integrin activation (Figure 4I).

Our results demonstrate that SIRPA localization is a key determinant of its activity. In the absence of CD47, SIRPA is relegated to the phosphatase-rich zone outside the cell bead interface (Freeman et al., 2016; Goodridge et al., 2011). This localization prevents SIRPA activation. Conversely, CD47 binding retains SIRPA at the Src-kinase rich phagocytic cup, where it is activated and suppresses engulfment. Spatial segregation of Src-family kinase activity at the central phagocytic synapse and CD45 phosphatase activity at the periphery underlies the activation of many activating receptors (TCR, Fc Receptor, (Freeman et al., 2016; James and Vale, 2012). Our work expands this model, suggesting that exclusion of inhibitory receptors like SIRPA may be a pre-requisite for efficient engulfment. Further, these data suggest a new paradigm for regulating inhibitory receptors based on conditional recruitment to the immunological synapse.

SIRPA exclusion from the phagocytic synapse in the absence of CD47 prevents basal inhibition of engulfment and allows positive signaling to dominate. This exclusion requires the extracellular domain of SIRPA, as replacing the extracellular domain with a small, inert protein (FRB) allowed SIRPA to enter the phagocytic synapse (Figure 2). CD45, the transmembrane phosphatase that negatively regulates Fc Receptor activation, is sterically excluded from the synapse between a T cell or macrophage and its target (Freeman et al., 2016; Goodridge et al., 2011; James and Vale, 2012). The SIRPA extracellular domain is predicted to be smaller than CD45 (aglycosylated proteins are 12 nm and 17 nm respectively (Chang et al., 2016; Hatherley et al., 2008). Biophysical studies have shown that proteins that are the same size or slightly smaller than the height of a cell-cell synapse are excluded from the synapse due to steric constraints (Schmid et al., 2016). Ligand binding is sufficient to drive synapse localization (Schmid et al., 2016). Thus SIRPA may be sterically excluded unless CD47 ligation overcomes the energetic barrier preventing SIRPA from entering the immunological synapse. Alternatively, other mechanisms, such as lateral crowding or interactions with the surrounding glycocalyx, could drive SIRPA exclusion from the synapse.

After addressing the mechanism of SIRPA activation, we sought to identify the targets of CD47-SIRPA signaling. Previous work has shown that SIRPA activation dramatically reduces global phosphotyrosine, including phosphorylation of mDia, paxillin, talin, alpha-actinin and non-muscle myosin IIA (Okazawa et al., 2005; Tsai and Discher, 2008). However, discerning between direct targets of SIRPA-bound phosphatases and indirect targets resulting from an upstream block in the engulfment signaling cascade has been challenging. Because blocking non-muscle myosin II decreases phagocytosis to a similar extent as CD47, myosin has been presumed to be the primary target of SIRPA (Chao et al., 2012; Tsai and Discher, 2008). However, we demonstrate that the inhibitory effect of CD47-SIRPA can be eliminated by re-activating integrin, suggesting that the direct targets of SIRPA-bound SHP phosphatases are upstream of integrin activation. SHP-2 has previously been shown to directly dephosphorylate Fak (Yu et al., 1998) and vinculin (Campbell et al., 2018), thus SHP-2 may act upon these key integrin regulators. However, given the broad specificity of SHP-1 and SHP-2, these phosphatases may dephosphorylate several targets at the phagocytic cup to suppress signaling.

Our work provides new insights into the connection between SIRPA and integrins. While phosphopaxillin (Tsai and Discher, 2008), has previously been shown to be affected by CD47-SIRPA, the relative importance of integrin signaling had not previously been addressed. We show that CD47-SIRPA prevents integrin activation, allowing macrophages to quickly discriminate between targets based on the presence of CD47. SIRPA overexpression has previously been shown to decrease surface levels of integrin over time (Liu et al., 2008). While this decrease in integrin expression does not explain how SIRPA immediately prevents phagocytosis of a CD47-bound target, it suggests that long term exposure to activated SIRPA may decrease overall phagocytic capacity, even of targets lacking CD47. In addition, SIRPA has been implicated in regulating cell motility, as fibroblasts lacking SIRPA have impaired motility (Alenghat et al., 2012; Inagaki et al., 2000; Motegi et al., 2003). In this case, un-ligated SIRPA may instead act downstream of integrin, as eliminating SIRPA decreases integrin responsiveness (Alenghat et al., 2012; Inagaki et al., 2000).

By suppressing integrin activation, CD47-SIRPA signaling may be able to suppress many different signaling pathways. Interestingly, CD47 has been reported to affect dendritic cell activation, cancer cell killing via a nibbling behavior (called trogocytosis), and complement-mediated engulfment (Caron et al., 2000; Matlung et al., 2018; Oldenborg et al., 2001; Tamada et al., 2004; Wu et al., 2018; Yi et al., 2015). These processes are triggered by diverse positive signaling receptors, but all require inside-out activation of integrin (Caron et al., 2000; Matlung et al., 2018; Oldenborg et al., 2001; Tamada et al., 2004; Wu et al., 2018; Yi et al., 2015). Targeting integrin, a common co-receptor, may explain how CD47-SIRPA signaling can regulate these diverse processes.

Finally, we found that integrin activation by manganese can drive engulfment of whole cancer cells by bone marrow derived macrophages. As a cancer treatment, CD47 blockade synergizes with therapeutic antibodies, like rituximab (Advani et al., 2018; Chao et al., 2010a). Activating integrins with a small molecule agonist in combination with antibody therapeutics may have a similar synergistic effect as CD47 blockade. Small molecule agonists of CD11b, an integrin subunit highly expressed in macrophages, drive tumor regression in a macrophage-dependent manner (Panni et al., 2019; Schmid et al., 2018). Our data suggests that these small molecules may allow macrophages to bypass the CD47 inhibitory signal.

## Acknowledgments

We thank K. McKinley and O. Klein for providing mouse long bones as a source for hematopoietic stem cells. We thank members of the Vale lab for critical feedback on this manuscript. M.A.M. was supported by the National Institute of General Medical Sciences of the National Institutes of Health under award number F32GM120990. This work was funded by the Howard Hughes Medical Institute.

## Materials and Methods

### Cell culture

RAW264.7 macrophages were provided by the ATCC and certified mycoplasma-free. The cells were cultured in DMEM (Gibco, Catalog #11965–092) supplemented with 1 × Pen-Strep-Glutamine (Corning, Catalog #30–009 Cl) 1 mM sodium pyruvate (Gibco, Catalog #11360-070) and 10% heat inactivated fetal bovine serum (Atlanta Biologicals, Catalog #S11150H). To keep variation to a minimum, cells were discarded after 20 passages. L1210 cells were also acquired from the ATCC.

J774A.1 macrophages were provided by the UCSF cell culture facility. J774A.1 and 293T cells were tested for mycoplasma using the Lonza MycoAlert Detection Kit (Lonza, Catalog# LT07-318) and control set (Lonza, Catalog #LT07-518).

Bone marrow derived macrophages were generated from the hips and long bones of C57BL/6J mice as previously described (Weischenfeldt and Porse, 2008) except that purified 25 ng/ml M-CSF (Peprotech, Catalog # 315–02) was used.

### Constructs and antibodies

All relevant information is provided in the STAR methods table, including a detailed description of the amino acid sequence of each construct and the catalog number of all antibodies.

### Lentivirus production and infection

All constructs were expressed in RAW264.7 using lentiviral infection. Lentivirus was produced in HEK293T cells transfected with pMD2.G (a gift from Didier Tronon, Addgene plasmid # 12259 containing the VSV-G envelope protein), pCMV-dR8.91 (since replaced by second generation compatible pCMV-dR8.2, Addgene plasmid #8455), and a lentiviral backbone vector containing the construct of interest (derived from pHRSIN-CSGW, see STAR methods) using lipofectamine LTX (Invitrogen, Catalog # 15338–100). Constructs are described in detail in the Key Resources Table. The media was harvested 72 hours post-infection, filtered through a 0.45 µm filter and concentrated using LentiX (Takara Biosciences). After addition of the concentrated virus, cells were centrifuged at 2000xg for 45 min at 37°C. Cells were analyzed a minimum of 60 hr later.

### Supported lipid bilayer assembly

#### SUV preparation

The following chloroform-suspended lipids were mixed and desiccated overnight to remove chloroform: 96.8% POPC (Avanti, Catalog # 850457), 2% Ni^2+^-DGS-NTA (Avanti, Catalog # 790404), 1% biotinyl cap PE (Avanti, Catalog # 870273), 0.1% PEG5000-PE (Avanti, Catalog # 880230, and 0.1% atto390-DOPE (ATTO-TEC GmbH, Catalog # AD 390–161). The lipid sheets were resuspended in PBS, pH7.2 (Gibco, Catalog # 20012050) and stored under argon. The lipids were broken into small unilamelar vesicles via several rounds of freeze-thaws. The mixture was cleared using ultracentrifugation (TLA120.1 rotor, 35,000 rpm / 53,227 × g, 35 min, 4°C). The lipids were then stored at 4°C under argon for up to two weeks.

#### Planar bilayer preparation for TIRF microscopy

Ibidi coverslips (catolog #10812) were RCA cleaned. Supported lipid bilayers were assembled in custom plasma cleaned PDMS (Dow Corning, catalog # 3097366-0516 and 3097358-1004) chambers at room temperature for 1 hour. Bilayers were blocked with 0.2% casein (Sigma, catalog # C5890) in PBS. Proteins were coupled to the bilayer for 45 min. Imaging was conducted in HEPES buffered saline (20 mM HEPEs, 135 mM NaCl, 4 mM KCl, 10 mM glucose, 1 mM CaCl_2_, 0.5 mM MgCl_2_). Bilayers were assessed for mobility by either photobleaching or monitoring the mobility of single particles.

#### Bead preparation

8.6* 10^8^ silica beads with a 5.02 µm diameter (10 µl of 10% solids, Bangs Labs, Catalog # SS05N) were washed three times with PBS, mixed with 1mM SUVs in PBS and incubated at room temperature for 0.5-2 hr with end-over-end mixing to allow for bilayer formation. Beads were then washed three times with PBS to remove excess SUVs and incubated in 100 µl of 0.2% casein (Sigma, catalog # C5890) in PBS for 15 min before protein coupling. Unless otherwise indicated, anti-biotin AlexaFluor647-IgG (Jackson ImmunoResearch Laboratories Catalog # 200-602-211, Lot # 137445) was added between 3 and 30 nM, always using the lowest IgG concentration that triggered engulfment. Purified CD47^ext^-His_10_ was added at 1 nm. Proteins were coupled to the bilayer for 1 hr at room temperature with end-over-end mixing.

#### Protein density estimation

Given the high affinity of His_10_ for Ni^2+^-DGS-NTA (0.6 nM (Hui and Vale, 2014)), and antibody-antigen interactions, we expect close to 100% coupling efficiency (Hui and Vale, 2014). Complete coupling would result in 600 molecules/µm^2^ CD47 and 300 molecules/µm^2^ IgG for the 3 nM coupling condition. This is well within the range of CD47 on the surface of a cancer cell (Figure S1). In addition, to estimate the amount of IgG bound to each bead, we compared the fluorescence of IgG on the bead surface to calibrated fluorescent beads (Quantum AlexaFluor 647, Bangs Lab) using confocal microscopy. Using this method, we measured 200-360 molecules/µm^2^ of IgG, which is consistent with the theoretical prediction of near complete coupling.

### Protein Purification

His_10_-CD47^ext^, His_10_-CD47^ext^ ^F37D, T115K^ (aa40-182; Uniprot Q61735) and ICAM-tagBFP-His_10_ (O’Donoghue et al., 2013) were expressed in SF9 or HiFive cells using the Bac-to-Bac baculovirus system as described previously (Hui and Vale, 2014). Briefly, the N-terminal extracellular domain of CD47 was cloned into a modified pFastBac HT A with an upstream signal peptide from chicken RPTPσ (Chang et al., 2016). Insect cell media containing secreted proteins was harvested 72 hr after infection with baculovirus. His_10_ proteins were purified by using Ni-NTA agarose (Qiagen, Catalog # 30230), followed by size exclusion chromatography using a Superdex 200 10/300 GL column (GE Healthcare, Catalog # 17517501). The purification buffer was 30 mM Hepes pH 7.4, 150 mM NaCl, 2 mM MgCl_2_, 5% glycerol (CD47) or 150 mM NaCl, 50 mM Hepes pH 7.4, 5% glycerol, 2 mM TCEP (ICAM).

### Phosphopaxilin staining

Macrophages were fixed in 4% PFA for 15 min, then permeabilized and blocked with 0.1% BSA in PBS with 0.5% Tween 20. The cells were incubated with the phosphopaxillin antibody at 1:50 dilution at 4° C overnight before incubating with Alexa Fluor 555 anti-rabbit secondary (21428), Alexa Fluor 488 phalloidin (A12379).

### β2 integrin block and Fab generation

To disrupt integrin function, the 2E6 anti-β2 integrin antibody (ThermoFisher, MA1805) or isotype control (ThermoFisher, 16-4888-81) was added to macrophages at 10 μg/ml 30 minutes before IgG-opsonized beads. To eliminate any effects of the Fc domain, we generated Fabs from these antibodies using the Pierce Fab separation kit (ThermoFisher 44985).

### Whole cell internalization assay

30,000 macrophages infected with GFP-CAAX were plated in a 96-well glass bottom MatriPlate (Brooks, Catalog # MGB096-1-2-LG-L). 2 hours prior to imaging, cells were washed into serum-free, phenol free DMEM for imaging. Manganese (SigmaAldrich, M8054) was added at 100 µM 30 min prior to imaging. or CD47 function-blocking antibody clone miap301 (Biolegend, 127520) was used at 10 mg/ml. 100,000 H2B-mCherry expressing L1210 cells were added and the co-culture was imaged for 8 hr.

### Microscopy and analysis

Images were acquired on a spinning disc confocal microscope (Nikon Ti-Eclipse inverted microscope with a Yokogawa spinning disk unit and an Andor iXon EM-CCD camera) equipped with a 40 × 0.95 NA air and a 100 × 1.49 NA oil immersion objective. The microscope was controlled using µManager. For TIRF imaging, images were acquired on the same microscope with a motorized TIRF arm, but using a Hamamatsu Flash 4.0 camera and the 100x 1.49 NA oil immersion objective.

### Quantification of engulfment

30,000 macrophages in one well of a 96-well glass bottom MatriPlate (Brooks, Catalog # MGB096-1-2-LG-L) between 12 and 24 hr prior to the experiment. Macrophages remained in culture media (DMEM with 10% heat inactivated serum) throughout the experiment. Unless otherwise indicated, ∼1 × 10^7^ beads were added to well and engulfment was allowed to proceed for 30 min. Cells were fixed with 4% PFA and stained with CellMask (ThermoFisher, catalog # C10045) without membrane permeabilization to label cell boundaries. Images were acquired using the High Content Screening (HCS) Site Generator plugin in µManager (Edelstein et al., 2010).

### Quantification of synapse intensity of phosphoPaxillin, actin and SIRPA constructs

Phagocytic cups were selected for analysis based on the presence of clustered IgG at the cup base (SIRPA chimeras) or clear initiation of membrane extensions around the phagocytic target (actin, phosphopaxillin). The phagocytic cup and the cell cortex were traced with a line 3 pixels wide at the Z-slice with the clearest cross section of the cup. The average background intensity was measured in an adjacent region and subtracted from each measurement.

### Quantification of the cell-bilayer contact area

For 3A, time-lapse images of macrophages interacting with an IgG or IgG+CD47 bilayer were acquired using TIRF microscopy as described above. Macrophages were removed from their culture dish using 5% EDTA in PBS, two times washed and resuspended in the HEPES imaging buffer (20 mM HEPEs, 135 mM NaCl, 4 mM KCl, 10 mM glucose, 1 mM CaCl_2_, 0.5 mM MgCl_2_) before being added to the TIRF chamber. The area of the cell contacting the bilayer was traced in ImageJ beginning with the first frame where the cell can be detected. Only cells with mobile IgG clusters were included. For 3B, the number of macrophage-bilayer contacts and the area was quantified in still images of live cells between 10 and 15 min after cells were added to the bilayer. All cells were included.

### Statistics

Statistical analysis was performed in Prism 8 (GraphPad, Inc). The statistical test used is indicated in the relevant figure legend.

## Supplemental Figure Legends

**Figure S1, related to Figure 1: A reconstitution system for studying CD47-SIRPA signaling**

(A) SDS page gel shows the N-terminal extracellular domain of murine CD47 purified from insect cells using a C-terminal His_10_. (B) Beads coated in supported lipid bilayers were incubated with the indicated concentration of anti-biotin IgG. The fluorescent intensity of Alexa Fluor 647-IgG on the bead was measured to ensure that the binding of IgG increased with higher coupling concentrations. (C) The estimated surface density of CD47 on red blood cells (Gardner et al., 1991; Mouro-Chanteloup et al., 2003), T cells (Subramanian et al., 2006), cancer cells (Dheilly et al., 2017; Jaiswal et al., 2009; Michaels et al., 2017) and the beads used in this study. (D) IgG surface density was held constant while CD47 density was titrated. The 1 nM CD47 coupling concentration was selected for use throughout this study. (E) Histograms depict the fraction of macrophages engulfing the indicated number of phosphatidylserine beads. The RAW264.7 histograms correspond to the replicates depictured in Figure 1E. RAW264.7 engulfment was measured after 30 min and J774A.1 was measured after 90 min.

**Figure S2, related to Figure 2: Forcing SIRPA into the macrophage-target synapse suppresses engulfment**

(A) Schematic depicts TIRF imaging. (B) TIRF microscopy of J774A.1 macrophages encountering a 10% phosphatidylserine bilayer reveals that SIRPA-GFP is depleted at the center off the cell-bilayer synapse (top; yellow arrow compared to cyan arrow). Macrophages did not form this zone of depletion when encountering a bilayer containing both phosphatidylserine and CD47 (bottom). The ratio of SIRPA-GFP fluorescent intensity at the cell center/cell edge is quantified on the right. Each dot represents an individual cell and data is pooled from 3 independent experiments. Lines denote the mean ± SD. * * * denotes p<0.0005 by Student’s T test. (C) SIRPA-GFP and the chimeric receptors FcRI^ext^-SIRPA^int^-GFP and FRB^ext^-SIRPA are expressed at similar levels. Fluorescent intensity was normalized to the average intensity of SIRPA-GFP in that experiment. Each dot represents an individual cell and data is pooled from 3 independent experiments. Lines denote the mean ± SD. (D, E and F) Histograms depict the fraction of macrophages engulfing the indicated number of IgG-bound beads. The average number of beads per cell is shown ± SEM. This data corresponds to 1E (D), 1F (E) and 1G (F). For all panels, data is pooled from three data is pooled from 3 independent experiments. Lines denote the mean ± SD.

**Figure S3, related to Figure 3: CD47 does not affect FcR activation and Syk recruitment.**

(A) TIRF microscopy shows that macrophages are able to form IgG microclusters (left; cyan in merged image) that recruit Syk (middle; magenta in merged image) if CD47 is absent (top) or present (bottom). Inset shows the boxed region of the image above. The linescan shows the fluorescent intensity of Alexa Fluor 647-IgG and Syk-mCherry at the indicated position (white arrow). Intensity was normalized so that 1 is the highest observed intensity and 0 is background. The fraction of cells able to form IgG clusters and recruit Syk is displayed on the far right. Each dot represents the percent from an independent experiment (n≥ 20 per replicate) and the lines denote mean ± SD. (B) TIRF microscopy shows that, in the presence of CD47, SIRPA (green) does not co-localize with IgG clusters (cyan; arrowheads). Inset shows the boxed region in the above image. The linescan shows the fluorescent intensity of Alexa Fluor 647-IgG and SIRPA-GFP at the position indicated by a white arrow. (C) Macrophages were incubated with a Fab generated from the β2 function-blocking antibody (2E6, red) or from an isotype control (green). The pooled data from three independent replicates is graphed with error bars denoting SEM. * * indicates p<0.005 by Kruskal-Wallis test.

**Figure S4, related to Figure 4: Manganese does not affect L1210 viability**

L1210 cells were serum starved for 2 hrs, then treated with 100 μM manganese for 6 hrs as in 2H. The percent of cells that bound high levels of annexin, indicating phosphatidylserine exposure and the initiation of apoptosis, was measured by flow cytometry.

**Movie S1: Macrophage encounters IgG bound to a supported lipid bilayer.**

TIRF imaging (see schematic in Figure S2) shows Alexa Fluor 647-IgG (black) in the supported lipid bilayer as a macrophage engages with an IgG-bound target. Frames were acquired every 20 sec and time is indicated in the top left. Scale bar denotes 5 µm.

**Movie S2: Macrophage encounters IgG and CD47 bound to a supported lipid bilayer.**

TIRF imaging shows Alexa Fluor 647-IgG (black) in the supported lipid bilayer as a macrophage engages with an IgG and CD47-bound target. Frames were acquired every 20 sec and time is indicated in the top left. Scale bar denotes 5 μm.

**Movie S3: An Mn-treated macrophage encounters an L1210 leukemia cell**

A macrophage infected with GFP-CAAX encounters an L1210 leukemia cell labeled with H2B-mCherry. In the presence of 100 μM Mn^2+^ the macrophage is able to engulf the cancer cell. Images were acquired every 5 min for 140 min. The field of view is 53 µm by 53 µm.

