## Supplementary figures and images for "CD47 suppresses phagocytosis by repositioning SIRPA and preventing integrin activation"

### Figure S1

#### Figure S1, related to Figure 1:
#### A reconsitution system for studying CD47-SIRPA signaling

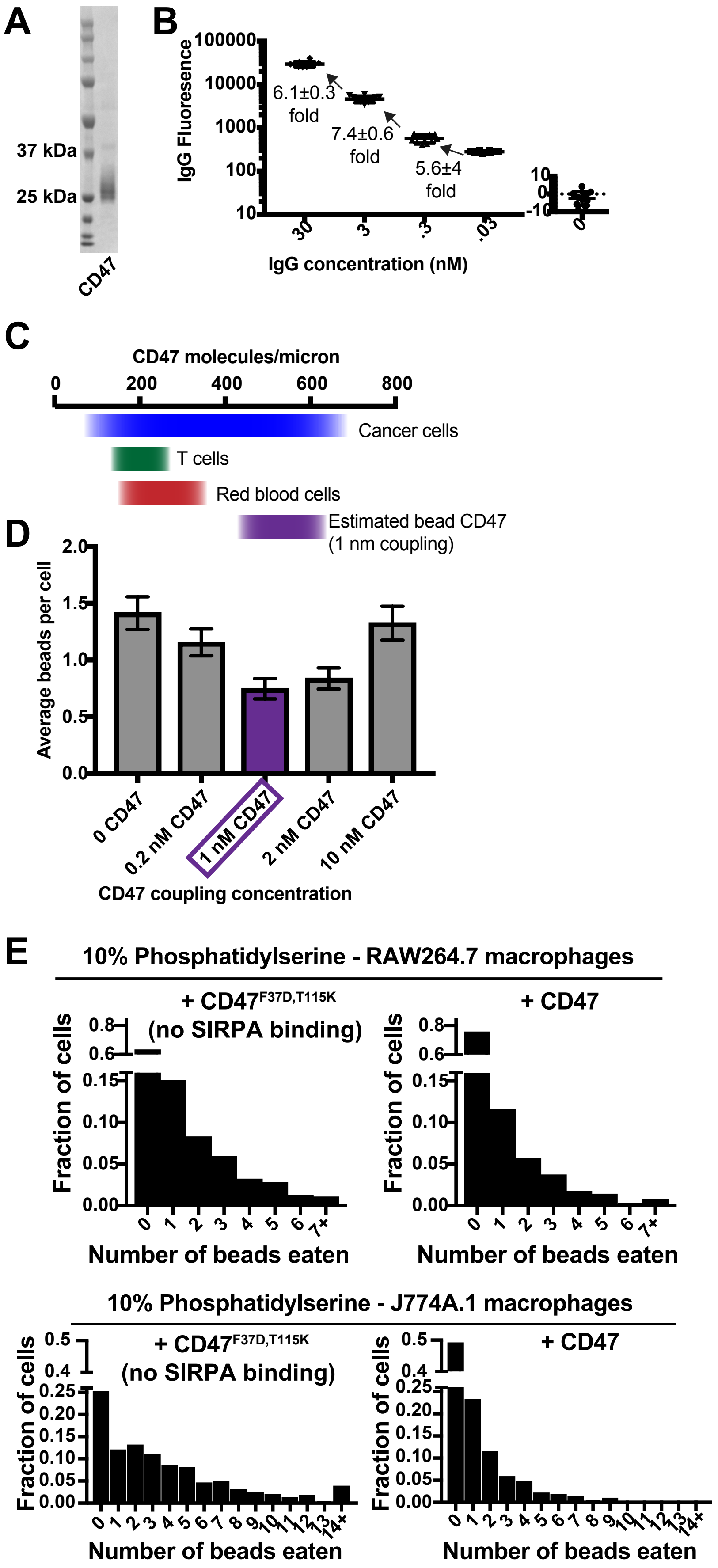

### Figure S3

#### Figure S3, related to Figure 3: CD47 does not affect Fc Receptor activation

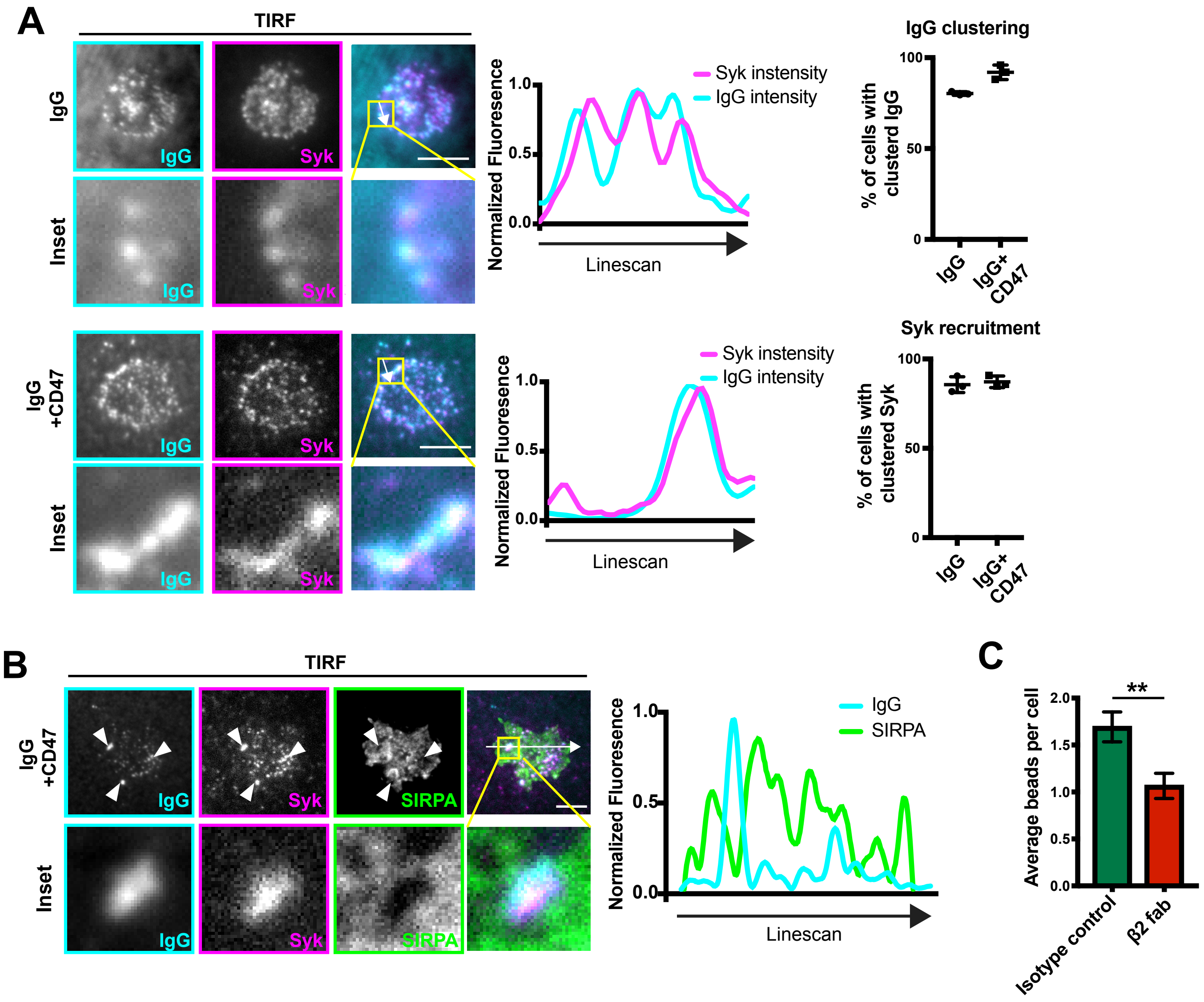

### Figure S4

#### Figure S4, related to Figure 4: Manganese does not affect L1210 viability

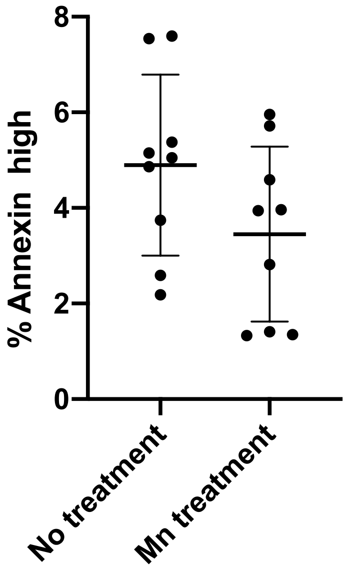
