## Supplementary material for "CD47 suppresses phagocytosis by repositioning SIRPA and preventing integrin activation": Figure S2

### Figure S2, related to Figure 2:
### Forcing SIRPA into the macophage-target synapse supresses engulfment

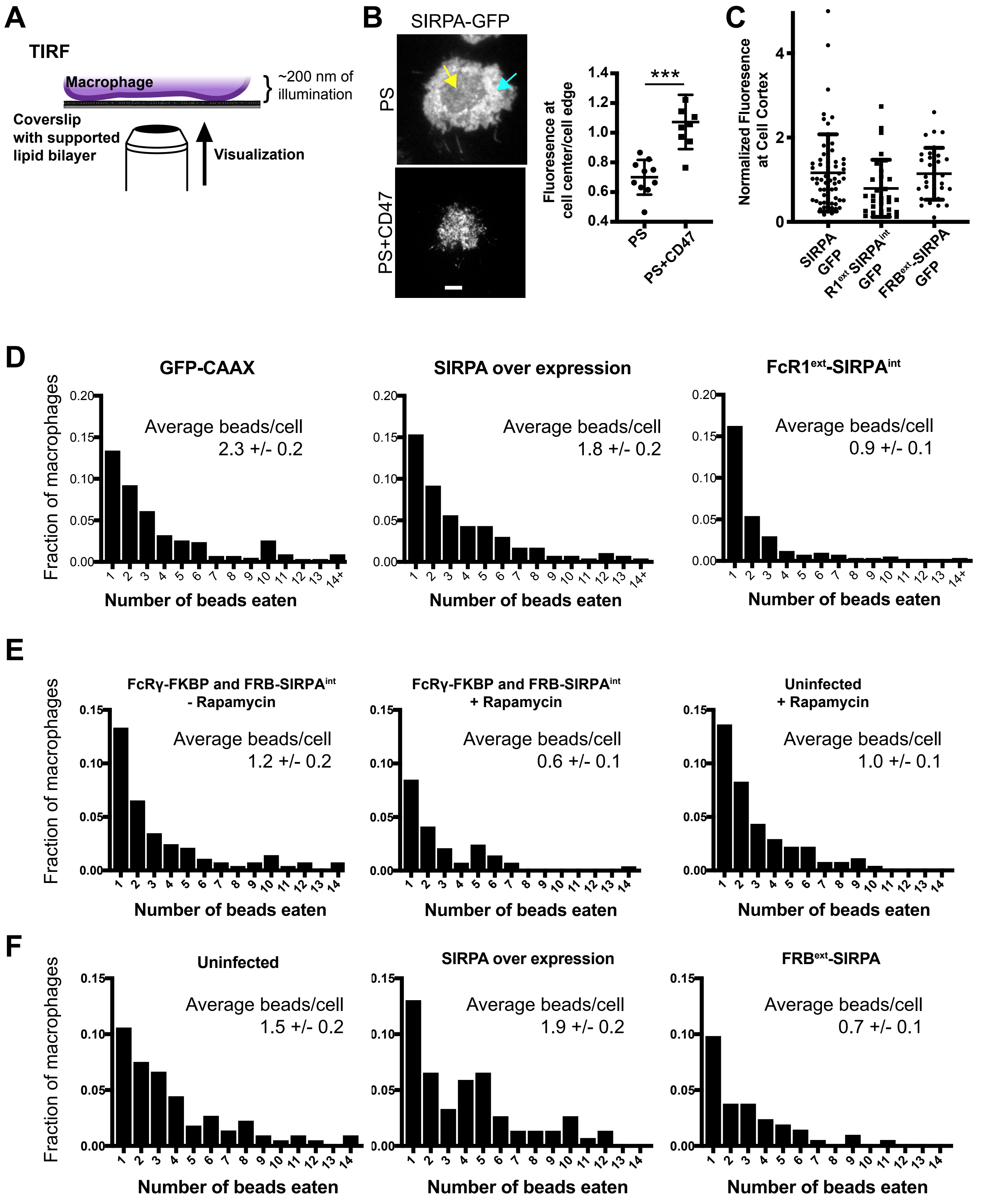
